# A NOVEL AND EFFICIENT ALGORITHM FOR DE NOVO DISCOVERY OF MUTATED DRIVER PATHWAYS IN CANCER

**DOI:** 10.1101/117473

**Authors:** By Binghui Liu, Chong Wu, Xiaotong Shen, Pan Wei

## Abstract

Next-generation sequencing studies on cancer somatic mutations have discovered that driver mutations tend to appear in most tumor samples, but they barely overlap in any single tumor sample, presumably because a single driver mutation can perturb the whole pathway. Based on the corresponding new concepts of coverage and mutual exclusivity, new methods can be designed for de novo discovery of mutated driver pathways in cancer. Since the computational problem is a combinatorial optimization with an objective function involving a discontinuous indicator function in high dimension, many existing optimization algorithms, such as a brute force enumeration, gradient descent and Newton's methods, are practically infeasible or directly inapplicable. We develop a new algorithm based on a novel formulation of the problem as non-convex programming and nonconvex regularization. The method is computationally more efficient, effective and scalable than existing Monte Carlo searching and several other algorithms, which have been applied to The Cancer Genome Atlas (TCGA) project. We also extend the new method for integrative analysis of both mutation and gene expression data. We demonstrate the promising performance of the new methods with applications to three cancer datasets to discover de novo mutated driver pathways.

## 1. Introduction

It is known that cancer is characterized by numerous somatic mutations, of which only a subset, named *’’driver”* mutations, contribute to tumor growth and progression. With next-generation whole-genome or whole-exome sequencing, somatic mutations are measured in large numbers of cancer samples (Mardis & Wilson, 2008;Meyerson *et al.,* 2010). To improve understanding and treatment of cancers, it is critical to distinguish driver mutations from neutral *”passenger”* mutations. A standard approach to predicting driver mutations is to identify recurrent mutations in cancer patients (*Beroukhim et al.,* 2007; *Getz et al., 2007*), which has its drawback in its inability to capture mutational heterogeneity of cancer genomes *(Ding et al., 2008*; Jones *et al.,* 2008). An emerging discovery is that in a given sample driver mutations typically target one, but not all, of several genes in cellular signaling and regulatory pathways (Vogelstein & Kinzler, 2004). Hence the research has shifted from the gene level to pathway level (Boca, 2010; Efroni, 2011). Recent studies indicated that mutations arising in driver pathways often cover a majority of samples, but, importantly, for a single sample only a single or few mutations appear because a single mutation is capable to perturb the whole pathway; the latter concept is the so-called *mutual exclusivity.* By using mutual exclusivity, new pathway-based methods are developed to identify de novo driver mutations and pathways *(Ciriello et al., 2012*; Masica *et al.,* 2011;Miller *et al.,* 2011). For example, Miller *et al.* (2011)proposed a method to find functional sets of mutations by using patterns of recurrent and mutually exclusive aberrations; Ciriello *et al*. (2012)not only used the mutual exclusivity pattern, but also incorporated a gene functional network constructed based on prior knowledge. Recently Vandin et *al.* (2012)introduced a novel scoring function combining the two concepts, **coverage** and **mutual exclusivity**, to identify mutated driver pathways through optimizing this scoring function, which has been used in some large-scale cancer sequencing studies. It is solved by stochastic search methods: a greedy algorithm and a Markov chain Monte Carlo method. Other proposals based on binary linear programming, genetic search algorithm, and integer linear programming have appeared (Zhao *et al.,* 2012; Leiserson *et al.,* 2013), all of which are still relatively slow, especially for large-scale problems. To address these issues, we reformulate the problem of identifying mutated driver pathways as a statistical problem of subset identification to minimize a new cost function, what we call minimum cost subset selection (MCSS). A key component is a novel approximation to a combinatorial problem through regularization, where a discontinuous indicator function is approximated by a continuous and non-convex truncated *L*_1_ (TL) function (Shen *et al.,* 2012). Furthermore, we add a truncated *L*_1_ penalty (TLP) to the cost function to seek a sparse solution, as well as adding a small ridge penalty to alleviate the problem of multiple solutions. As a result, a combinatorial optimization problem becomes a continuous but non-convex one in the Euclidean space, which can be efficiently solved through a non-convex optimization technique, leading to high computational improvement.

Another advantage of the proposed method is that it is able to find multiple mutated driver pathways. An existing method to identify multiple mutated driver pathways is Multi-Dendrix (Leiserson *et al.,* 2013), in which the number of pathways and the number of the genes in each pathway have to be specified in advance. On the contrary, our proposed method does not need to fix such numbers beforehand. Based on a series of randomly selected initial estimates, a series of low-cost estimates of mutated driver pathways can be obtained. Moreover, the proposed method is general so that other types of information can be incorporated in a simple way. For example, if a gene interaction network is available, it can be incorporated by adding a network-based penalty to the current cost function as in Li &Li (2008); since it is more informative to combine mutation data with other types of data such as gene expression data (Zhang and Zhou, 2014), an integrative version can be developed by adding other cost functions for other types of data into the current one. As a concrete example, we propose a new method to integrate mutation data with gene expression data.

## 2. Methods

### 2.1. Problem

Consider <u>mutation data</u> with *n* patients and *p* genes, represented as an *n* × *p* mutation matrix ***A*** with entry *A_ij_* = 1 if gene *j* is mutated in patient *i*, and *A_ij_* = 0 otherwise. For gene *j* ∈ *V* = {1,…,*p*}, let Γ (*j*) = {*i*: *A_ij_* = 1} be a subgroup of patients whose gene *j* is mutated. Moreover, given a subset of genes *B* ⊆ {1, …,*p*}, let Γ(*Β*) be a subgroup of patients with at least one of the genes in *B* mutated, i.e. Γ(*Β*) = ∪_*j*∈*B*_ Γ(*j*). Cancer sequencing studies have motivated to identify a set of mutated genes across a large number of patients, whereas only a small number of patients have mutations in more than one gene in the set, that is, these mutations are approximately exclusive. This amounts to finding a set *B* ⊆ *V* of genes such that (i) the coverage is high, that is, most patients have at least one mutation in *B*; (ii) the genes in *B* are approximately exclusive, that is, most patients have no more than one mutation in B. A measure *ω*(*Β*) = ∑ _*j*∈*B*_ |Γ(*j*)| – |Γ(*Β*)| was proposed by Vandin *et al.* (2012), called the coverage overlap, to balance the trade-off between coverage and exclusivity. To maximize the coverage |Γ(*Β*)| and minimize the coverage overlap *ω*(*Β*) simultaneously, Vandin et *al.* (2012)suggests to minimize

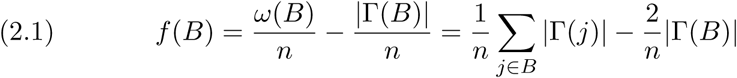

with respect to *Β*, thus obtaining an estimate *Β̂.* Minimizing *f* (*Β*) is equivalent to maximizing the weight function –*f* (*Β*), which is called the maximum weight sub-matrix problem (MWSP). Note that minimizing *f* (*Β*) is a non-trivial combinatorial problem, to which most existing optimization algorithms based on the gradient descent or Newton’s algorithm cannot be directly applied. A popular method called <u>Dendrix</u> is based on a Monte Carlo search algorithm to seek an approximate solution to minimize *f* (*B*) (Vandin *et al.,* 2012).

### 2.2. New formulation

As indicated in (2.1), MWSP is a combinatorial problem, for which a brute force search is time-consuming and not scalable for large (*n*,*p*), while many existing algorithms like gradient descent or Newton’s method cannot be directly applied. Here we formulate it as nonconvex minimization and examine a regularized version by imposing penalties to ensure proper solutions. Specifically, for any *β* e ∈^*p*^, let *B* = *B*(*β*) = {*j* ∈ *V*: *|*β*_j_|* ≠ 0}, and we rewrite 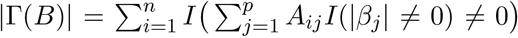, 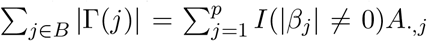, 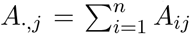 and 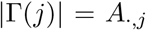 for each *j* ∈ {1,⋯,*p*}. Then (2.1) becomes

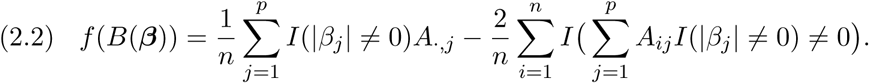

Minimizing (2.2) in *β* yields an estimate *β̆* = (*β̆*_1_,⋯, *β̆p*)′, and thus an estimated set *B̆* = {j: *| β̆*|= 0}. However, due to the discontinuity with the indicator function *I*(.), it is difficult to minimize (2.2) directly; instead, since min(|*β_j_|/τ*_1_,1) → *I*(*|β_j_|* ≠ 0) as *τ*_1_ *→* 0^+^, we propose a surrogate to minimize

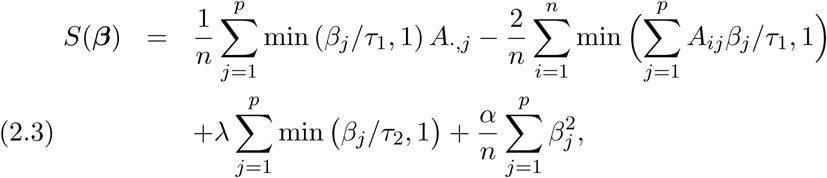

with respect to *β* = (*β*_1_,…, *β_p_*)*′* ∈ [0, +∞)^*p*^; that is, *β* is a vector of parameters to be estimated; λ, *α, τ*_1_ and *τ*_2_ are non-negative tuning parameters to be determined via a grid search in cross-validation (as used in the later experiments); *A_ij_*’s are observed and known. Note that in (2.3), the last two terms, as a TLP and a ridge penalty respectively, ensure sparse and proper solutions.

### 2.3. Computation

To solve nonconvex minimization (2.3), we employ difference convex (DC) programming by decomposing the objective function into a difference of two convex functions, on which convex relaxation is performed through iterative approximations of the trailing convex function through majorization. Specifically, 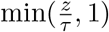 can be written as a difference of two convex functions: min(*z/τ*, 1) = *z/τ —* max [*z/τ —* 1, 0) for any *z* > 0 and *τ >* 0. Then, we obtain a sequence of upper approximations *S*^(*m*)^(*β*) of *S*(*β*) at iteration *m* (up to a constant) as follows:

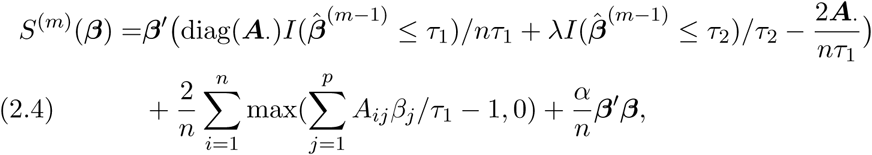

where *β* ∈ [0, +∞)^*p*^, ***A***.⋅= (*A*._,1_,*…, A*._,p_)′, and diag(***A***.) is a diagonal matrix with elements of ***A***. as diagonals. Now *S*^(*m*)^(*β*) is strictly convex (since the first term is linear in *β*, the second is convex while the last is quadratic in *β* with *α* ≠ 0), we use some existing convex program package (CVX in Matlab), or more efficiently, the subgradient descent method (as shown in the appendix) (Shor, 1985), to obtain a unique minimizer *β̂*^(m)^; we repeat the process until convergence to obtain *β̃* = *β̂*^(+∞)^.

Interestingly, one may replace the TLP in *S*(*β*) in (2.3) with the *L*_1_*-* penalty, yielding *β̂^L^*. This, together with, other randomly generated numbers, can be use as an initial value *β̂*^(0)^ for our method. For selection of tuning parameters, we may consider cross-validation, as discussed later. The following algorithm summarizes our computational method.

#### ALGORITHM 1

*Given the parameters τ*_1_, *τ*_2_, λ, *α.*

**Initialization** *supply an initial estimate β̂*^(0)^

**Iteration** *At iteration m, compute β̂*^(*m*)^ *by minimizing* (2.4).

**Stopping rule** *Terminate when S*(*β̂*^(*m*–1)^) – *S(β̂*^(*m*)^) ≤ 0. *The estimate is β̃* = *β̃*^(*m*\*–1)^ *, where m*\* *is the smallest index satisfying the termination criterion. The estimated subset is B̃* = {*j* ∈ {1,⋯*,p}*: *β̃_j_* = 0}.

The following convergence property of Algorithm 1 has been established.

#### THEOREM 1.

*β̂*^(*m*)^ *in Algorithm 1 converges in finite steps to a local minimizer β̃* of *S*(*β*) *in* (2.3)*. S*(*β̂*^(*m*)^) *strictly decreases in m until β̂*^(*m*)^ *= β̂*^(*m–*1)^ = *β̂*^(*m*–*1)^ *for all m* ≥ *m*\*.

### 2.4. Initial estimate

In general, a large number of good or randomly selected initial estimates may be used to obtain multiple solutions, from which a subset of more promising ones with smaller objective or cost function values can be selected. Below, we describe a simple way to obtain a good initial estimate, which was used in later simulations; we modify *S*(*β*) such that the modified version *S_L_*(*β*) becomes much easier to optimize. A local condition of (2.3) can be established based on regular subdifferentials

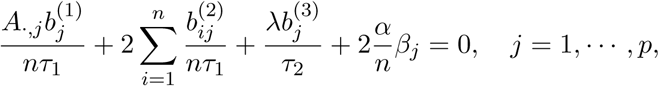

where 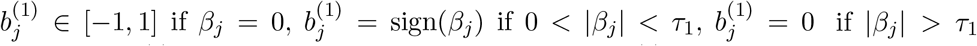 and 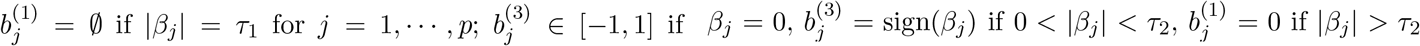 and 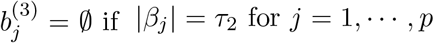 Note that 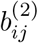 is more complicated as it depends on the values of *A_ij_*_′_ and 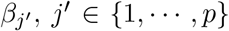, and 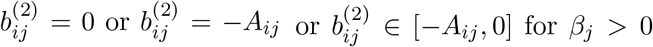. Based on these regular subdifferentials, we develop the following lemma.

#### LEMMA 1.

*If there exists a non-zero local minimizer β* of S*(*β*) *in* (2.3) *on* ℝ^*p*^, *then* 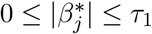 *for each j* ∈ {1,⋯,*p*}.

Lemma 1 says that the set of all local minimizers of *S*(*β*) in (2.3) over [0, +∞)]^*p*^ is the same as that obtained from the following cost function over [0,*τ*_1_]^*p*^:

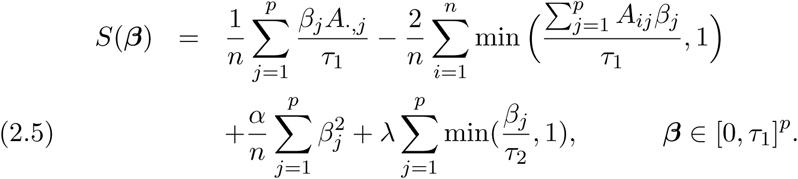

If we use the *L*_1_-penalty as opposed to the truncated *L*_1_-penalty in (2.5), then the cost function becomes

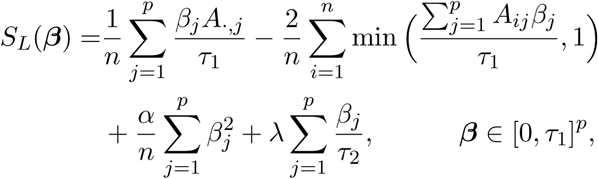

which is strictly convex in β ∈ [0,*τ*_1_]^*p*^, yielding a unique minimizer *β̂_L_.*

### 2.5. Model selection

Tuning parameters (λ, *r*) need to be estimated from data, where *τ*_2_ = *rτ*_1_ (0 < *r* < 1), while *α* is fixed at a sufficiently small positive number, say *α* = 10^−3^, and *τ*_1_ is fixed at any positive value, say *τ*_1_ = 1. Tuning of (λ, *r*) can be achieved through sample splitting. As a matter of fact, the term 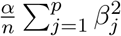 is introduced to yield a unique minimizer of (2.4) so that the bias caused by the ridge penalty is ignorable for sufficiently small *α*. On the other hand, given the ratio *r*, an exact value of (*τ*_1_,*τ*_2_) is unimportant. This is because 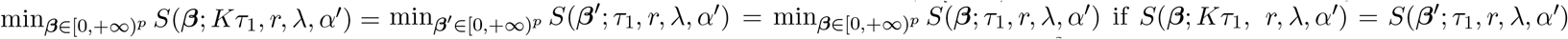 with *β*' = *β*/*Κ* and 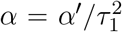 for any *K* > 0. Consequently, given the ratio *r*, optimization in terms of different choices of *τ*_1_ are equivalent. Given a *n* × p mutation matrix ***A***, a candidate set Λ ⊆ (0, +∞) of the tuning parameter λ and a candidate set *R* ⊆ (0, +∞) of the tuning parameter *r* = *τ*_2_/*τ*_1_, we use a sample splitting procedure to select the tuning parameters λ*̂* ∈ Λ and *r̂* ∈ *R:*

**Initialization** supply a randomly selected initial estimate *β̂*^(0)^.

**Partition** Randomly partition the rows of the mutation matrix *A* into two parts: training data ***A***^*tr*^ and tuning data ***A***^*tu*^.

**Training** For each λ ∈ Λ and each *r* ∈ *R*, apply Algorithm 1 to the training data *A^tr^* with the initial estimate *β̂*^(0)^ and parameters λ and *r* to get the corresponding estimate *β̂^tr^* (λ, *r*).

**Tuning** Based on the tuning data *A^tu^*, we formulate a tuning error for each *β̂^tr^* (λ, *r*) as

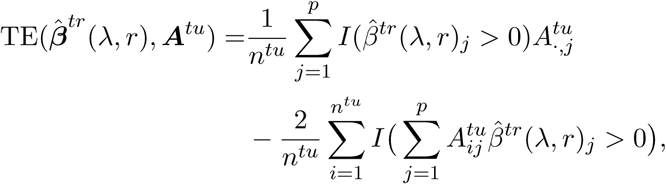

where *n^tu^* denotes the number of rows of *A^tu^*, that is, the patient number in the tuning data, and 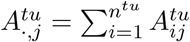. We select λ and *r* as

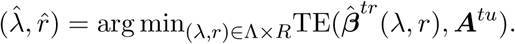

Given λ = λ*̂* and *r* = *r̂*, we apply Algorithm 1 to the original mutation matrix ***A*** to find *β̂* ∈ [0, +∞)^*p*^ that minimizes *S*(*β*) in (2.3).

### 2.6. Integrative analysis

An advantage of the proposed algorithm is its possible extensions to include other types of genomic data, in addition to mutation data. To this end, we modify the proposed cost function and algorithm to incorporate other types of data such as gene expression. Let *f_ME_*(*B*) denote the integrative cost function, which is the sum of the original cost function *f* (*B*) and a new one *f_E_*(*B*) for gene expression data:

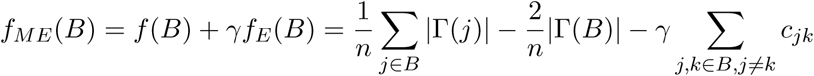

where *c_jk_* is the Pearson correlation coefficient of the expression profiles of genes *j* and *k*. Note that the integrative cost function is based on the observation that the genes in the same pathway usually collaborate with each other to execute a common function. Therefore, the expression profiles of the genes in the same pathway usually have higher correlations than those from different pathways (Qiu e*t al.,* 2010; Zhao et *al.,* 2012).

To minimize *f_ME_*(*B*), we develop a similar algorithm as before, called MCSS_ME, where *S*(*β*) and *S*^(*m*)^(*β*) are replaced by *S_ME_*(*β*) and 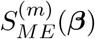 respectively as follows.

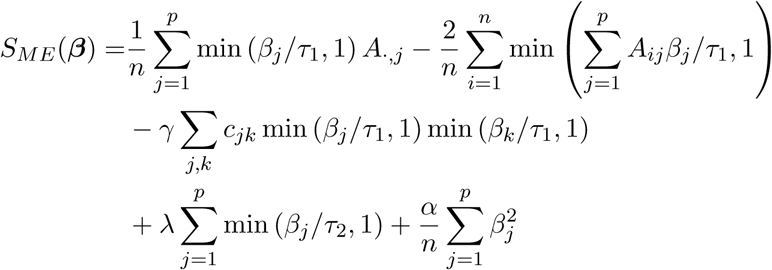

and

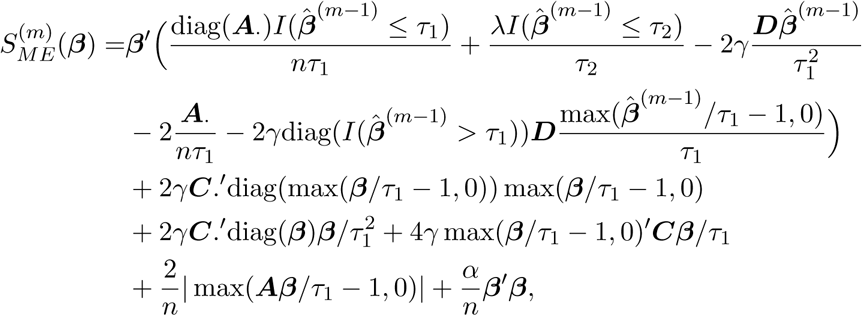

where ***D*** = ***C***+diag(***C***.), **C** = [*c_jk_*] (*c_jj_* = 0) and ***C***. is the row sum vector of ***C***. Here we use the subgradient descent method (as shown in the appendix) to obtain a minimizer of 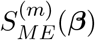.

To choose a suitable γ in situations with no prior information, we propose a method to balance the contributions to the new cost function from mutation data and from gene expression data. Specifically, we randomly select a large number of subsets, say *B*_1_, B_2_,…, *B_R_*, of the genes from {1, 2,⋯,*p*} with the size of each subset |*B_j_*| randomly generated from {2,3, ⋯ *,n_p_*}, then we choose γ = min_*j*_ f (*B_j_*)/min_*j*_ *f_E_*(*B_j_*), which aims to give an equal weight on the contribution of the mutation data and that of the expression data to the overall cost function *f_ME_*(). In our following experiments, we always used *R* = 10000 and *n_p_* = 8, though other values may be used.

After determining *γ*, we choose the other tuning parameters similarly as before but according to an integrative version of the tuning error

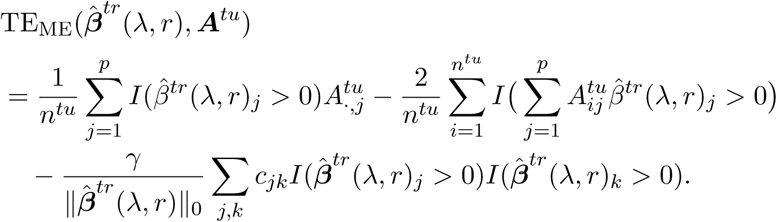

#### 2.6.1. Evaluation metrics

Several metrics are used for evaluation, including the correct (C) or incorrect (IC) numbers of non-zero estimates for the mutations/genes in the true pathway *B*_0_, and average differences of the cost function values (ADC) between the true set *B*_0_ and the estimated set B of the driver mutations/genes; that is, C=|*B*_0_ ∩ *B̂*|, 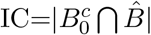, ADC= (*f*(*B*_0_) – *f*(*B̂*))/*n*. We also included the runnng time (RT) (in minutes) of each algorithm. Note that ADC is important, because the basic task for minimum cost subset selection is to identify a set of mutations with the minimum cost.

In addition to using the correct (C) or incorrect (IC) numbers of non-zero estimates and ADC to measure how close the estimated pathways are close to the true pathway, we also investigate several other metrics in decomposing the cost function into the coverage (*c_c_*) and exclusivity (*c_e_*), and displaying the proportion of the patients carrying a mutation of a gene in a pathway (*c*_1_), as well as the proportion of those carrying multiple mutations in more than one gene in the pathway (*c*_2_). Specifically, we define

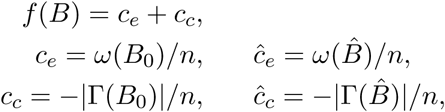

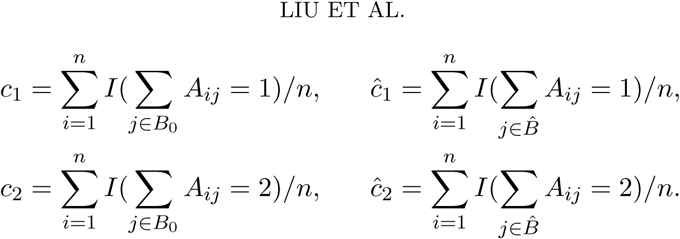

Due to the coverage and exclusivity of a pathway, *c*_1_ is often similar to — *c_c_* while *c*_2_ is similar to *c_e_.*

## 3. Results

### 3.1. Real data examples

In this section we first illustrate the application of the proposed method to two cancer datasets that were previously examined by Vandin et *al.* (2012), then to a more recent and larger dataset including both mutation and expression data. As argued by Vandin *et al.* (2012), a set of mutated genes with a low cost function value is likely to be a mutated driver pathway, based on which our primary objective is to identify such mutated driver pathways through minimum cost subset selection of mutated genes. For each of the first two datasets, the proposed method was applied with the tuning parameter λ chosen from a tuning set of size 10, while 100 randomly generated initial estimates were used. For each initial estimate, we applied the proposed method, by which we identified multiple low-cost sets of mutations.

#### 3.1.1. Lung adenocarcinoma

The original data set contains 1013 somatic mutations in 623 sequenced genes from 188 lung adenocarcinoma patients in the Tumor Sequencing Project (Ding *et al.,* 2008). For our purpose, we examined 356 genes that were mutated for at least one patient from a group of 162 patients, as in Vandin *et al.* (2012).

The proposed method was applied to identify multiple sets of mutated genes with low cost function values. Using 100 randomly selected initial values for MCSS, it cost 0.85 minutes and identified some gene sets with low cost. To demonstrate the resulting low-cost sets of mutations as possible candidates for mutated driver pathways, in Table 1 we group these discovered sets in terms of known pathways. In Table 1, all the discovered sets related to two known pathways associated with lung adenocarcinoma: the *mTOR* signaling pathway and the cell cycle pathway. Gene interactions in these pathways were reported in Ding *et al*. (2008)as depicted in Figure 1.

**TABLE 1.**
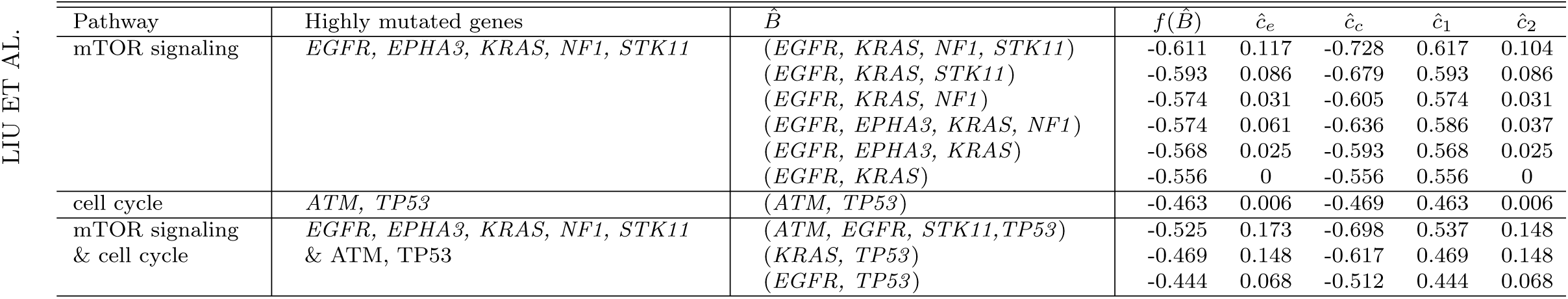
Applied to the mutation data of lung adenocarcinoma *(Ding et al, 2008),* the new method MCSS identified multiple sets of low-cost mutated genes, grouped in terms of associated pathways.

**FIG 1.**
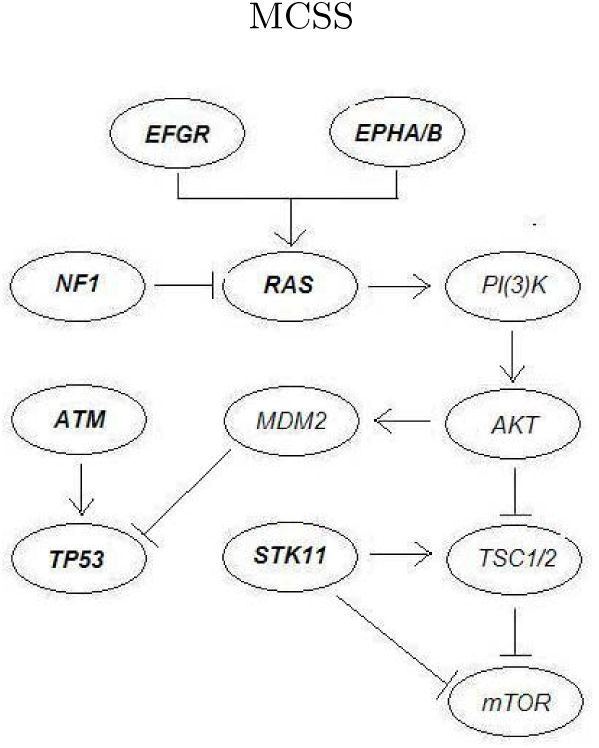
The mTOR signaling pathway and the cell cycle pathway associated with lung adenocarcinoma as reported in Ding et al. (2008). The KRAS gene is one of the three oncogenes in the Ras family.

First, as indicated in Figure 1 (see Figure 6 of Ding *et al.* (2008)), the *mTOR* signaling pathway consists of some highly mutated genes, such as *EGFR, EPHA3, KRAS, NF1* and *STK11. EGFR* is a well-known oncogene, whose mutations are strongly associated with lung cancer (da Cunha Santos *et al.,* 2011). In contrast, *EPHA3* is one of the most frequently mutated genes in lung cancer, which however has not yet been extensively investigated. As suggested by Zhuang et *al.* (2012), tumor-suppressive effects of wild-type *EPHA3* could be overridden in trans by dominant negative *EPHA3* somatic mutations discovered in patients with lung cancer. *KRAS* is an oncogene associated with non-squamous non-small cell lung cancer. As indicated by many studies as well as our analysis, the mutations of *KRAS* and *EGFR* are strongly mutually exclusive. *KRAS* serves as a mediator between extracellular ligand binding and intracellular transduction of signals from the *EGFR* to the nucleus. The presence of activating KRAS mutations has been identified as a potent predictor of resistance to EGFR-directed antibodies (Heinemann *et al.,* 2009). *STK11* encodes a tumor suppressor enzyme, and its mutations can allow cells to grow and divide uncontrollably, leading to the formation of cancerous cells (Gill *et al.,* 2007). In particular, *STK11* mutations are found in non-squamous non-small cell lung cancer, however uncommon in most other types of cancer.

Interestingly, all the identified sets of mutated genes with the cost function values *f* (*B̂*) lower than −0.556 = 90/162 are related to these five genes. Recall that inDing *et al.* (2008), *(EGFR, KRAS*) (*f* (*B̂*) = –0.556) and (*KRAS, STK11*) (*f* (*B̂*) = –0.420) are the most significant pairs in the mutual exclusiveness test, and in Vandin *et al.* (2012), the triplet (*EGFR, KRAS, STK11*) (*f*(*B̂*) = –0.593) was found with a lower cost, which was reported as a novel discovery. As indicated in Table 1, we could find not only this triplet (the second set in Table 1), but also another set (*EGFR, KRAS, NF1, STK11*) (*f* (*B̂*) = –0.611) (the first set in Table 1) that contains this triplet and has a lower cost function value. It is a better characterized gene set, containing the already discovered (*EGFR, KRAS, STK11*). In addition, we also identified four low-cost sets: *(EGFR, KRAS, NF1*) (*f* (*B̂*) = –0.574), (*EGFR, EPHA3, KRAS, NF1*) (*f* (*B̂*) = –0.574), (*EGFR, EPHA3, KRAS*) (*f*(*B̂*) = –0.568) and (*EGFR, KRAS*) (*f*(*B̂*) = –0.556). These discoveries suggest possible roles of these genes related to the mTOR signaling pathway.

Second, the cell cycle pathway includes two highly mutated genes, *ATM* and *TP53. ATM* plays a central role in cell division and DNA repair, and the protein encoded by this gene is an important cell cycle checkpoint kinase, which functions as a regulator of a wide variety of downstream proteins. Some studies suggested that *ATM* mutations may increase the risk for lung cancer (Lo *et al.,* 2010). On the other hand, *TP53* encodes a tumor suppressor protein p53 that regulates cell division by keeping cells from growing and dividing too fast or in an uncontrolled way. *TP53* mutations are the most common genetic changes found in human cancer, in particular as one of the most significant events in lung cancer while playing an important role in the tumorigenesis of lung epithelial cells (Ding *et al.,* 2008).

The pair (*ATM, TP53*) was identified by the proposed method with the cost function value of −0.463, which was also discovered in Vandin *et al.* (2012)by removing the triplet (*EGFR, KRAS, STK11*) from the original dataset. Note that among the identified low-cost sets in Table 1, the cost function value of (*ATM, TP53*) was relatively high due to its low value of the coverage: |Γ(*Β̂*)| = 76, much smaller than the maximum value of *n* = 162. As hypothesized inVandin *et al.* (2012), the low coverage is possibly because somatic mutations were measured in only a small subset of genes, or because only single-nucleotide mutations and small indels in these genes were measured, and other types of genomic or epigenetic alterations might occur in the ”unmutated” patients.

In addition, we identified some low-cost sets consisting of the genes related to both the mTOR signaling and the cell cycle pathways, namely, (*ATM, EGFR, STK11, TP53*) (*f* (*Β̂*) = –0.525), (*KRAS, TP53*) (*f* (*Β̂*) = –0.469) and (*EGFR, TP53*) (*f*(*Β̂*) = –0.444). Presumably these discoveries are related to that *EGFR* and *KRAS* are upstream regulators of *TP53,* as suggested by Ding *et al.* (2008).

#### 3.1.2. Glioblastoma multiforme (A)

Next, we analyzed the mutation data of 84 glioblastoma multiforme (GBM) patients from The Cancer Genome Atlas (The Cancer Genome Atlas Research Network, 2008), where 601 somatic mutations in these patients occurred. The mutation data consist of 84 patients and 178 genes, with each mutation occurring in at least one patient. The proposed method was applied to identify multiple sets of mutations with low cost values. Using 100 randomly selected initial values for MCSS, it cost 0.66 minutes and identified some gene sets with low cost. In Table 2 we also group the identified low-cost sets in terms of the possibly associated pathways. Most of the sets are associated with three important pathways of glioblastoma multiforme: the p53 signalling pathway, the RB signalling pathway and the RAS/RTK/PI(3)K signalling pathway. Interactions in these pathways were reported in The Cancer Genome Atlas Research Network (2008) as described in Figure 2. Below we discuss each pathway and the discovered sets of mutations.

**FIG 2.**
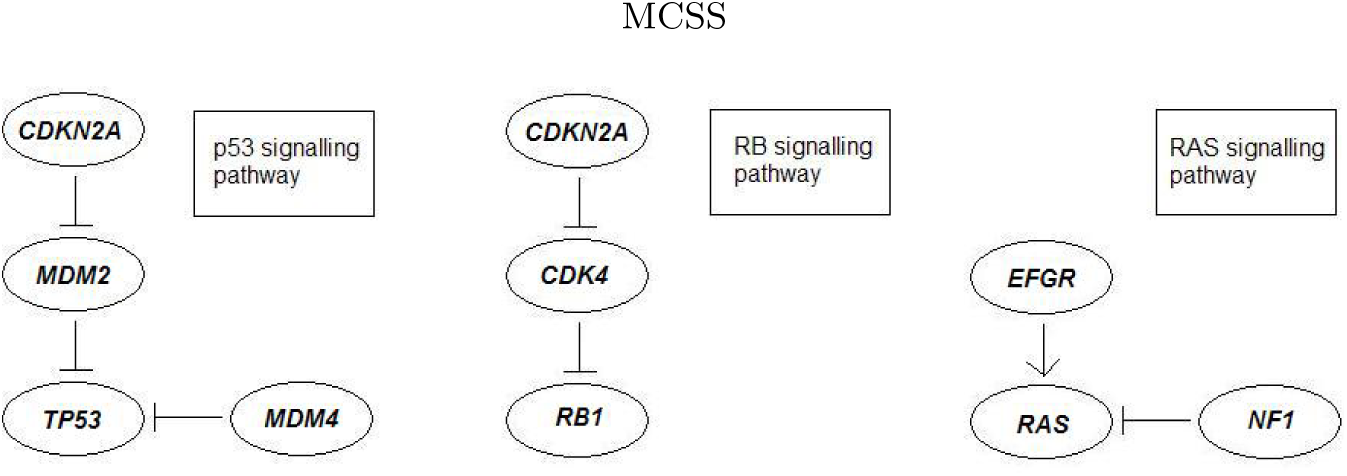
Three pathways associated with glioblastoma multiforme as reported in The Cancer-Genome Atlas Research Network (2008).glioblastoma multiforme.

**TABLE 2.**
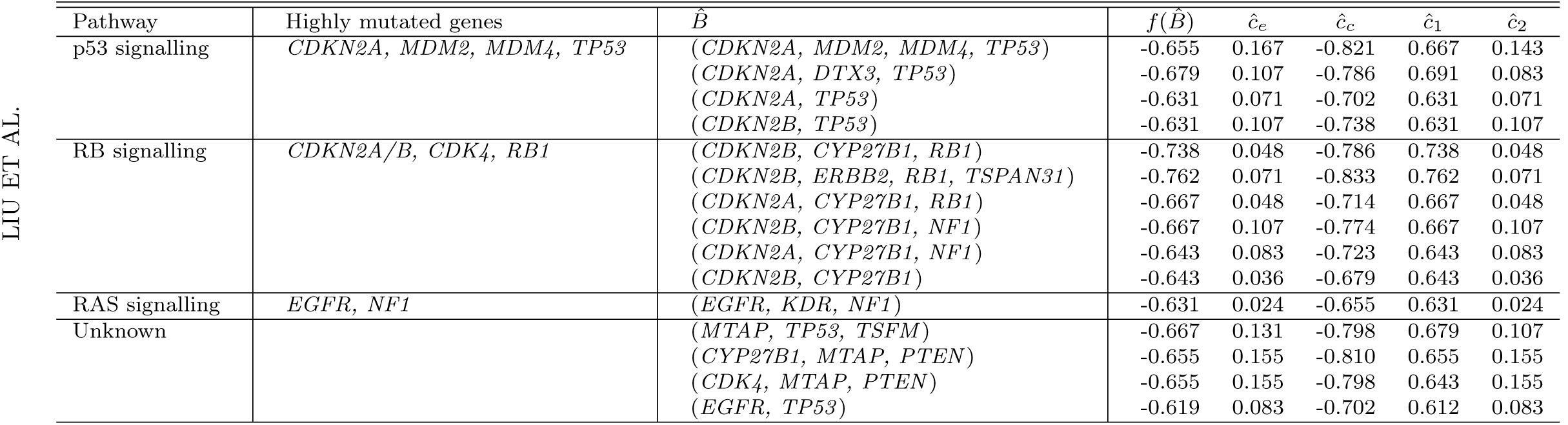
Applied to the mutation data of glioblastoma multiforme (data CBM A) (The Cancer Genome Atlas Research Network, 2008), the new method MCSS identified multiple sets of low-cost mutated genes, grouped in terms of associated pathways.

First, the p53 signalling pathway consists of some highly mutated genes, *CDKN2A*, *MDM2*, *MDM4* and *TP53.* Importantly, mutations in the tumour suppressor gene *TP53* are typical events in primary glioblastoma multiforme, which is characterised by a short clinical history and the absence of a pre-existing, less malignant astrocytoma. In contrast, the cellular oncogene MDM2 is viewed as an important negative regulator of the p53 tumor suppressor, whose overexpression is a characteristic feature of secondary glioblastoma multiforme, progressing from less malignant astrocytoma (Stark *et al.,* 2003).

Interestingly, the set of these four genes (*CDKN2A, MDM2, MDM4, TP53*) (*f* (*Β̂*) = −0.655 = −55/84) was identified by the proposed method as a novel discovery unreported before, e.g., in comparison with the pair (*CDKN2A, TP53*) (*f*(*Β̂*) = −0.631) identified by Vandin *et al.* (2012). As indicated in Table 2, the pair (*CDKN2A, TP53*) was also uncovered by the proposed method, in addition to another two sets, (*CDKN2A, DTX3, TP53*) (*f*(*Β̂*) = −0.679) and (*CDKN2B, TP53*) (*f*(*Β̂*) = –0.631). Since *CDKN2A* and *CDKN2B* are tumor suppressor genes located on a common homozygous deletion region on the human genome, they mutate almost simultaneously, which leads to a low cost function value of (*CDKN2B, TP53*). However, for (*CDKN2A, DTX3, TP53*), currently without further biological evidence, we conjecture that it has a low cost function value mainly because it consists of a low-cost set (*CDKN2A, TP53*) and gene *DTX3* with infrequent mutations.

Second, the RB signalling pathway consists of some highly mutated genes, *CDKN2A/B, CDK4, RB1,* where *CDKN2A* and *CDKN2B* are tumor suppressor genes, whose gene products, p16INK4A and p15INK4B, are both able to inhibit the binding of *CDK4* and *CDK6* to cyclin D, preventing the cell cycle progression at G1 phase. As a result, by negatively controlling cell cycle progression, these genes function as a critical defense against tumorigenesis of a great variety of human cancers, including glioblastoma multiforme (Feng *et al.,* 2012). The main set of mutations identified by the proposed method and associated with this pathway is likely to be (*CDKN2B, CYP27B1, RB1*) (*f*(*B̂*) = –0.738) since it has very low cost and often overlaps with other sets with low cost, which is coincided with that identified by Vandin et *al.* (2012). Since the mutational profile of *CYP27B1* is nearly identical to a metagene including *CDK4,* Vandin *et al.* (2012)believed that the triplet (*CDKN2B, CDK4, RB1*) may be of interest. For (*CDKN2B, CYP27B1, RB1*), the low cost function value is mainly due to the inclusion of *CDKN2B* and *CYP27B1.* As shown in Table 2, we identified several other sets containing *CDKN2A/CDKN2B* and *CYP27B1,* namely, (*CDKN2A, CYP27B1, RB1*) (*f*(*B̂*) = –0.667), (*CDKN2B, CYP27B1, NF1*) (*f*(*B̂*) = –0.667), *(CDKN2A, CYP27B1, NF1*) (*f*(*B̂*) = –0.643) and (*CDKN2B, CYP27B1*) (*f*(*B̂*) = –0.643). In addition, we also uncovered a set *(CDKN2B, ERBB2, RB1, TSPAN31*) (*f*(*B̂*) = –0.762), which is another new discovery by the proposed method. Interestingly, *TSPAN31* belongs to the same metagene including *CDK4.*

Third, the RAS/RTK/PI(3)K signalling pathway consists of some highly mutated genes, *EGFR, NF1, PI(3)K* and *PTEN.* Associated with this pathway, we identified a set of (*EGFR, KDR, NF1*) (*f*(*B̂*) = –0.619). Its low cost function value is likely due to the inclusion of *EGFR* and *NF1.*

Finally, among the other identified low-cost sets in Table 2, (*MTAP, TP53, TSFM*) (*f*(*B̂*) = −0.667), (*CYP27B1, MTAP, PTEN*) (*f*(*B̂*) = –0.655) and (*CDK4, MTAP, PTEN*) (*f*(*B̂*) = –0.655) are not known to be related to the pathways associated with glioblastoma multiforme. Hopefully, these low-cost sets will be useful for suggesting new links to glioblastoma multiforme. For (*EGFR, TP53*) (*f*(*B̂*) = –0.619), its low cost function value is possibly due to the approximate exclusiveness of *EGFR* and *TP53.* In particular, tumors in the ‘classical’ subtype of glioblastoma multiforme often carry extra copies of *EGFR* and are rarely mutated in *TP53.*

In summary, as shown in the above two real data examples, nearly all of the identified low-cost sets by the proposed method are associated with some known mutated driver pathways. This suggests potential usefulness of the proposed method. More importantly, in comparison with an existing method, some new discoveries were obtained, such as (*EGFR, KRAS, NF1, STK11*) (*f*(*B*) = –0.611 = –99/162) associated with the mTOR signalling pathway of lung cancer, and (*CDKN2A, MDM2, MDM4, TP53*) (*f* (*B̂*) = –0.656 = –55/84) associated with the p53 signalling pathway of glioblastoma multiforme.

#### 3.1.3. Glioblastoma multiforme (B)

Finally we analyzed a larger dataset of glioblastoma multiforme (GBM) patients from The Cancer Genome Atlas (Brennan *et al.,* 2013). The mutation data consist of 291 patients and 9539 genes, while the gene expression data include 558 patients and 12042 genes. Focusing on the intersection of the two gene sets, we obtained 5959 genes. Hence, we studied the filtered mutation data with 291 patients and 5959 genes, and the filtered expression data with 558 patients and 5959 genes.

First, the proposed MCSS was applied to identify multiple sets of mutations with low cost values using only the filtered mutation data. Using 10000 randomly selected initial values for all genes and 10000 randomly selected initial values for the subset of the genes with mutation rate larger than 0.05, MCSS identified some top gene sets with the six lowest cost function values (Table 3); note some gene sets with tied cost function values. They are mainly the variations and combinations of two core sets, (EGFR, KEL, NF1, TP53) and (IDH1, PIK3CA, PTEN), as contained in the top two sets identified. The list includes many well-known GBM genes, such as EGFR, PTEN, IDH1, TP53 and NF1 (Frattini *et al.,* 2013). Nevertheless, it is surprising that some top genes identified in Table 2 do not show up in the current list. Accordingly, we examined the top gene sets identified in the previous section but calculated their cost function values using the current data. From Table 4, we see that the top sets obtained earlier all have higher (i.e. worse) cost function values than those obtained in Table 3, indicating some inherent differences between the two datasets. For example, some high-mutation genes in the previous dataset, such as CDKN2A, MDM2, MDM4, CDKN2B, CYP27B1, ERBB2 and TSPAN31, had a low-mutation rate <5% in the current dataset. We use the less frequent mutation (LFM) (i.e. with a mutation rate < 5% among the subjects) ratio (i.e. the proportion of the LFM genes in a gene set) to indicate the presence of LFM genes in Table 4. The inherent differences between the two datasets confirm the genomic heterogeneity of GBM, one of the biggest challenges in current data analysis.

**TABLE 3.**
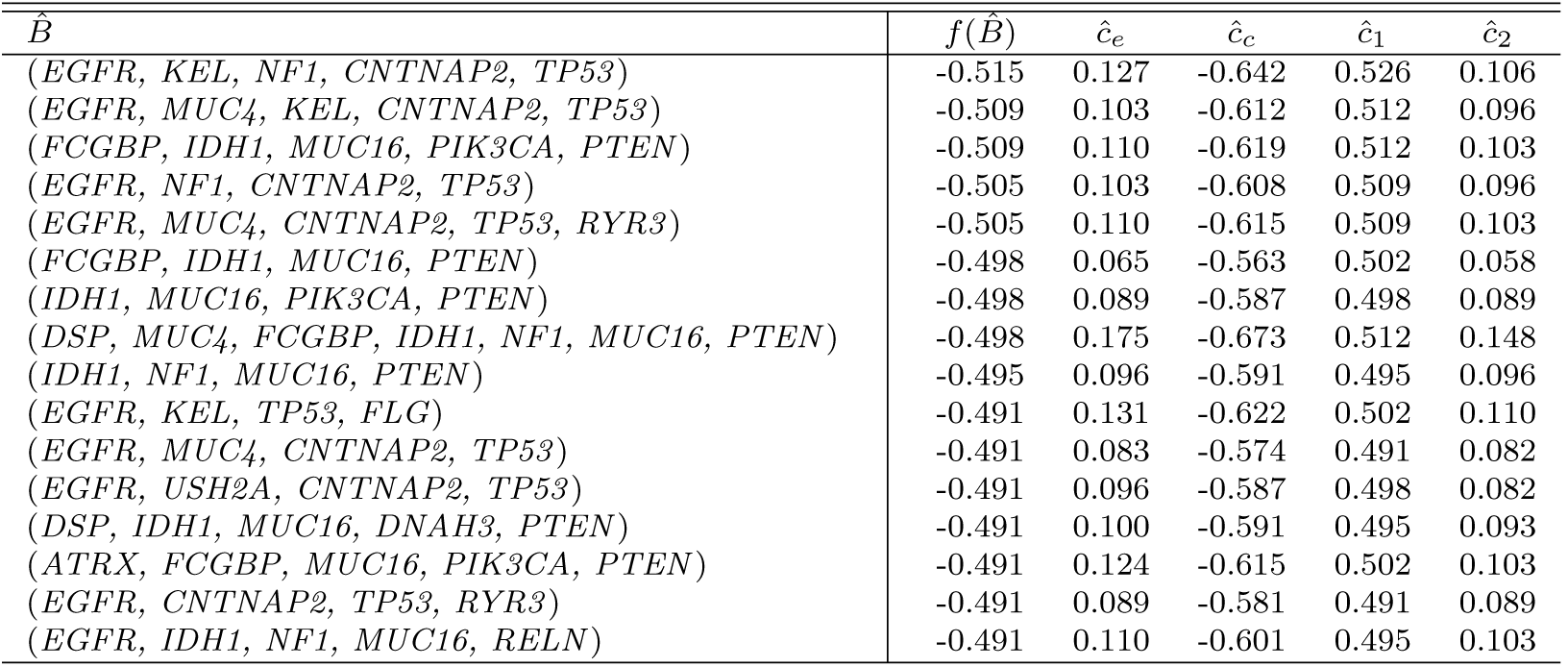
Application to the mutation data of glioblastoma multiforme (data GBM B) (Brennan et al., 2013): the top gene sets with the six lowest cost function values identified by the new method MCSS.

**TABLE 4.** The cost function values of the gene sets in the larger GBM (B) dataset with the gene sets identified from the smaller GBM (A) dataset.

| $\hat{B}$ | $f(\hat{B})$ | $\hat{c}_e$ | $\hat{c}_c$ | $\hat{c}_1$ | $\hat{c}_2$ | LFM ratio |
| --- | --- | --- | --- | --- | --- | --- |
| (CDKN2A, MDM2, MDM4, TP53) | -0.285 | 0.007 | -0.292 | 0.285 | 0.007 | 3/4 |
| (CDKN2A, TP53) | -0.289 | 0 | -0.289 | 0.289 | 0 | 1/2 |
| (CDKN2B, TP53) | -0.285 | 0 | -0.285 | 0.285 | 0 | 1/2 |
| (CDKN2B, CYP27B1, RB1) | -0.103 | 0 | -0.103 | 0.103 | 0 | 2/3 |
| (CDKN2B, ERBB2, RB1, TSPAN31) | -0.103 | 0 | -0.103 | 0.103 | 0 | 3/4 |
| (CDKN2A, CYP27B1, RB1) | -0.107 | 0 | -0.107 | 0.107 | 0 | 2/3 |
| (CDKN2B, CYP27B1, NF1) | -0.124 | 0 | -0.124 | 0.124 | 0 | 2/3 |
| (CDKN2A, CYP27B1, NF1) | -0.124 | 0 | -0.124 | 0.124 | 0 | 2/3 |
| (CDKN2B, CYP27B1) | -0.127 | 0 | -0.127 | 0.127 | 0 | 2/2 |
| (EGFR, KDR, NF1) | -0.354 | 0.024 | -0.378 | 0.354 | 0.024 | 1/3 |
| (EGFR, TP53) | -0.447 | 0.048 | -0.495 | 0.447 | 0.048 | 0/2 |

Finally, MCSS_ME was applied in an integrative analysis of both the filtered mutation and gene expression data. We did not apply the GA method because its current implementation requires the same set of the subjects with both mutation and gene expression data, which did not hold here. Using 10000 randomly selected initial values, MCSS_ME identified its top 10 gene sets shown in Table 5. We note that several genes were also identified from the other dataset in the previous section. Many selected genes are annotated in the Cancer Gene Census in the Catalogue Of Somatic Mutations In Cancer (COSMIC) (Forbes *et al.,* 2015), including well-known GBM genes (EGFR, PTEN, IDH1, TP53 and NF1, among others) (Frattini *et al.,* 2013). Here we only highlight a few examples. Gene ATRX was an important member of the H3.3-ATRX-DAXX chromatin remodelling pathway, among the most frequently mutated genes in paediatric and adult GBM Schwartzen-truber et *al.* (2012). Gene PIK3CA, encoding a protein that antagonizes the function of PTEN protein in the PI3K/Akt pathway; an exclusive mutation pattern was observed in PIK3CA and PTEN (Hartmann et *al.*, 2005). Mutations in a single gene, IDH1, resulted in reorganization of the methylome and transcriptome in glioblastomas and other cancers (Turcan *et al.,* 2012). As reviewed in Sturm et *al.* (2014), unsupervised clustering of the gene expression data from 200 adult GBM samples from TCGA identified four different molecular subtypes: proneural, neural, classical and mesenchymal. The proneural subtype was largely characterized by abnormalities in platelet derived growth factor receptor *α* (PDGFRA) or isocitrate dehydrogenase 1 (IDH1), whereas mutation of the epidermal growth factor receptor (EGFR) was found in the classical subgroup and mutations in neurofibromin (NF1) were common in mesenchymal tumours. In particular, Sturm et *al.* (2014)mentioned the detection of lower-frequency events in both cancer-related as well as previously un-associated genes such as ATRX and KEL.

**TABLE 5.**
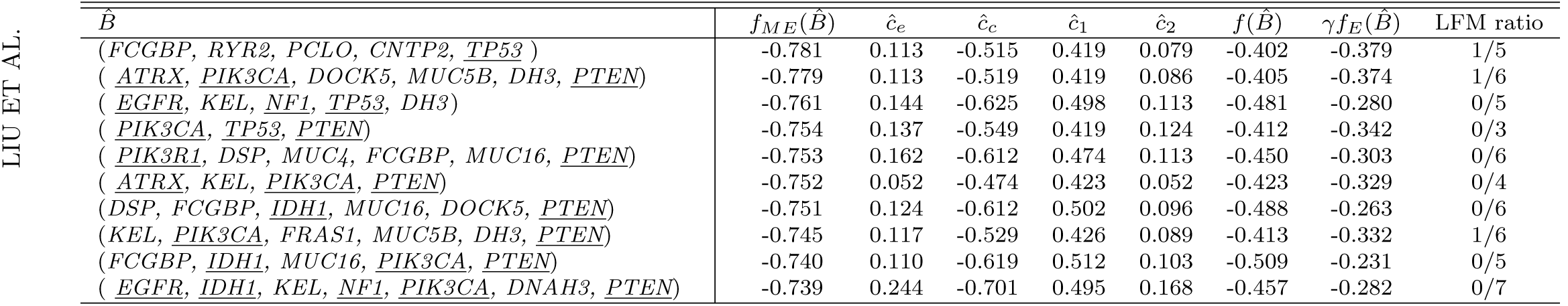
Application to the mutation data and gene expression data of glioblastoma multiforme (Brennan et al, 2013): the top 10 gene sets identified by the new method MCSS-ME with the automatically selected *γ* = 0.1. The known cancer genes annotated on COSMIC are underlined.

Note that all the gene sets identified with only the mutation data include only high-mutation genes (i.e. with a mutation rate > 5% among the subjects), while it is of interest but difficult to identify driver genes with less frequent mutations (i.e. with a mutation rate ≤ 5%) (LFM). Hence, we show the LFM ratio in Table 5. It is interesting to note the presence of two LFM genes, CNTP2 and DH3. In summary, our preliminary results seem to support the use of integrative analysis as advocated by others (Frattini *et al.,* 2013).

### 3.2. Simulations

Due to the difficulties in evaluating de novo discoveries with real data, we performed extensive simulations to study the operating characteristics of the proposed method and compared its performance against its competitors. All simulations were performed on a single processor of an Intel(R) Xeon(R) 2.83GHz PC.

#### 3.2.1. Simulation I: a single driver pathway

We first considered the case with only a single driver pathway, in which the focus was on comparing our new method with its strong competitor, the MCMC algorithm of Dendrix as implemented in Python (Vandin *et al.,* 2012), though several other methods ere also included.

For the proposed method, we fixed τ_1_ = 1, τ_2_ = 0.1 and α = 10^−3^, and tuned λ over a tuning set Λ. Specifically, λ was selected by minimizing a tuning error over a set of 10 equally-spaced points. We used 100 random initial estimates for MCSS (based on the subgradient descent algorithm), containing the Lasso estimate *β̂_L_*, as well as the other 99 random initial estimates. For the algorithm of Dendrix Vandin et *al.* (2012), 1000000 iterations were run for MCMC with sampling sets of size 4 for every 1000 iterations. Moreover, the algorithm was run with the number of driver mutations varying from 1 to 10 to select the best fitted subset with the lowest cost of *f* (⋅) in (2.1) as the final result.

For each simulated dataset, an *n* × *p* mutation matrix ***A*** was generated with a 1 indicating a mutation and 0 otherwise. For each patient, a gene in a driver pathway *B*_0_ = {1,2,3,4} was randomly selected and it mutated with probability *p*_1_, and another gene in *B*_0_ was randomly selected to have a mutation with probability *p*_2_. Consequently, *p*_1_ and *p*_2_ controlled the coverage and exclusiveness of *B_0_* respectively. Other genes outside *B*_0_ mutated with probability *p*_3_. Six set-ups were examined with (*p*_1_,*p*_2_,*p*_3_) = (0.95, 0.01, 0.05): (1) *n* = 50 and *p* = 1000, (2) *n* = 100 and p = 1000, (3) *n* = 1000 and *p* = 50, (4) *n* = 1000 and *p* = 100, (5) *n* = 50 and *p* = 10000, (6) *n* = 100 and *p* = 10000. With (*p*_1_,*p*_2_,*p*_3_) = (0.8,0.02,0.05), we had similar set-ups. The simulation results are summarized in Tables 6 and 7.

**TABLE 6.**
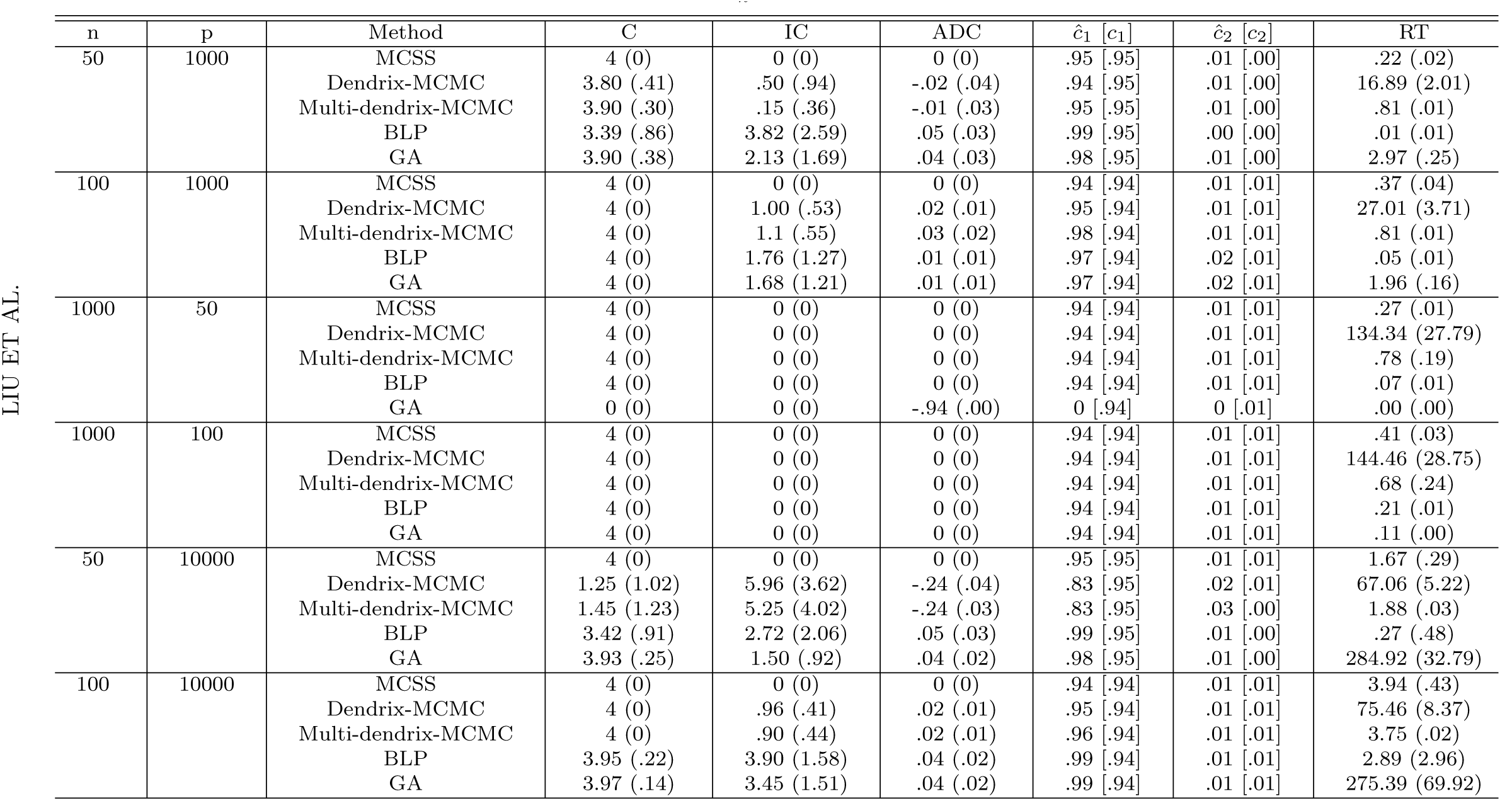
Results in Simulation I based on 100 simulation replications with (*p*_1_,*p*_2_,*p*_3_) = (0.95,0.01,0.05). The sample means (SD in parentheses) of correct (C) or incorrect (IC) numbers of non-zero estimates, average differences of the cost (ADC) between the true gene subset B_0_ = {1,2, 3,4} and the estimated subset B, that is, 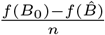,and the running time (RT) (in minutes) of the algorithms.

**TABLE 7.**
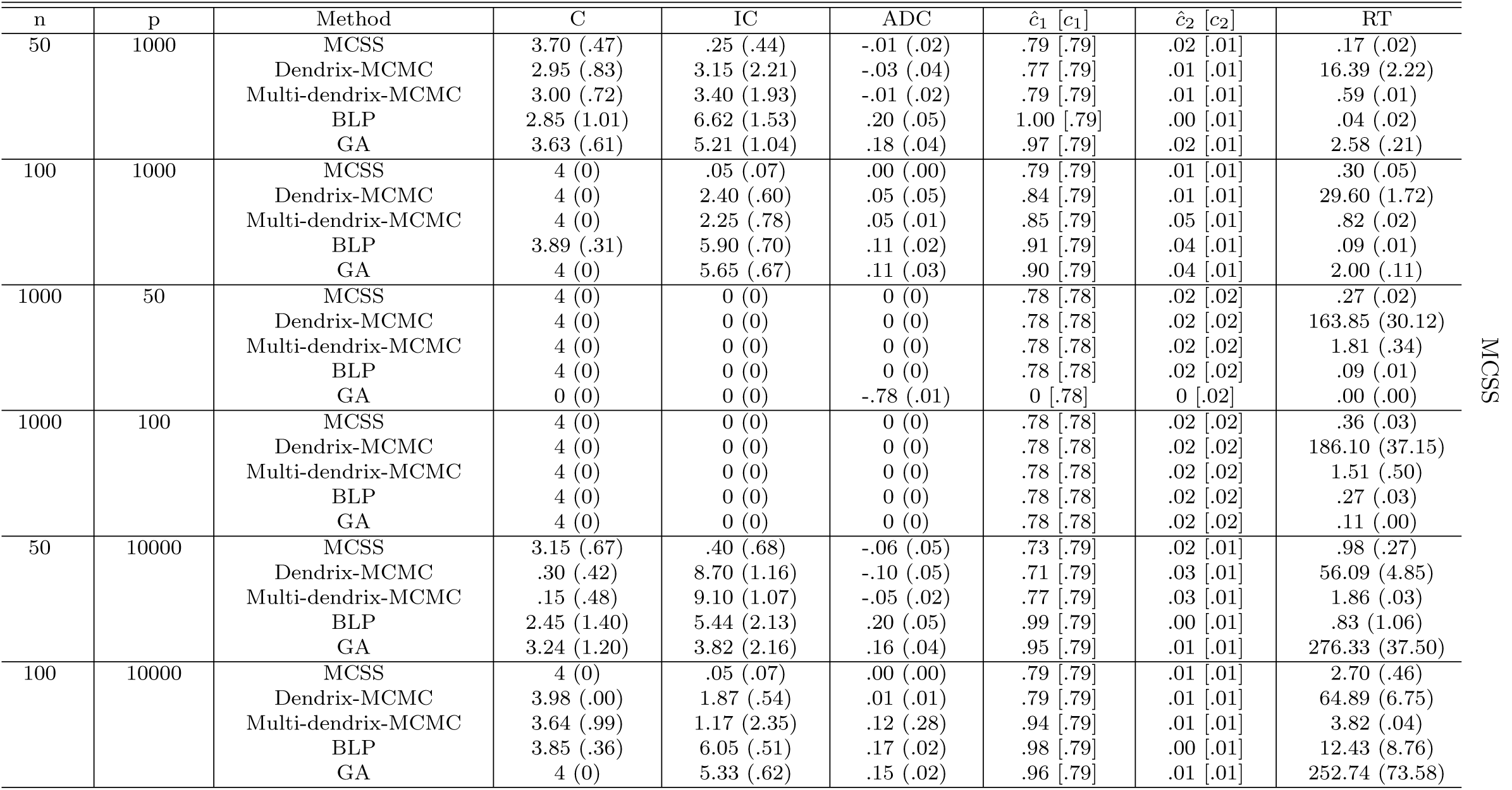
Results in Simulation I based on 100 simulation replications with (*p*_1_,*P_2_*,P_3_) =(0.8,0.02,0.05).

As suggested in Tables 6 and 7, the proposed method outperformed the MCMC algorithm of Dendrix, especially in the high-dimensional situations, with respect to the accuracy of selection as well as computational efficiency measured by the values of C, IC, ADC and RT respectively. The amount of improvement of the proposed method over the competitor ranged from low to high. For the running time, the proposed algorithm was overwhelmingly faster than the MCMC algorithm of Dendrix. In particular, it was often more than 50 times faster than the MCMC algorithm of Dendrix. As expected, both methods tended to perform worse as the amount of coverage and exclusiveness of a mutated driver pathway decreased.

We also compared our new method with several other alternative methods that were proposed more recently, including Multi-dendrix-MCMC ofLeiserson et *al.* (2013), BLP (binary linear programming) and GA (genetic algorithm) ofZhao et *al.* (2012). The numerical results of the three methods are also summarized in Tables 6 and 7. These results suggest that the performance of Multi-dendrix-MCMC was quite similar to that of Dendrix-MCMC but much faster; BLP and GA performed better than their competitors if the algorithms could finish running; however, they were not robust with frequent running errors (up to 15% failing to converge or giving output properly). In particular, BLP ran quite unsteadily in high-dimensional situations, say *n* = 50 and *p* = 1000 or 10000, while GA was too slow in high-dimensional situations since it tried to seek an exact solution. As expected, we see that these three methods also tended to perform worse as the amount of coverage and exclusiveness of a mutated driver pathway decreased. Since a rarely mutated gene may by chance satisfy the (approximate) exclusivity property with a highly mutated gene, the union of the highly mutated gene and some rarely mutated genes could drive down the cost function value, leading to false positives. To investigate this issue, we conducted a simulation study. As before, the driver pathway contained four genes. We set the 1st gene to have mutation in a fraction 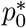 of all *n* patients, while setting the other three driver genes {2, 3, 4} to have mutations only in the remaining patients, for whom a gene from {2, 3, 4} was randomly selected with probability 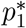 to have a mutation, and another gene in {2,3,4} was randomly selected to have a mutation with probability 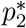. Finally, other genes outside *B*_0_ = {1,2,3,4} mutated with a background probability 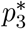. The corresponding simulation results are summarized in Table 8, suggesting that the proposed method still performed well.

**TABLE 8.**
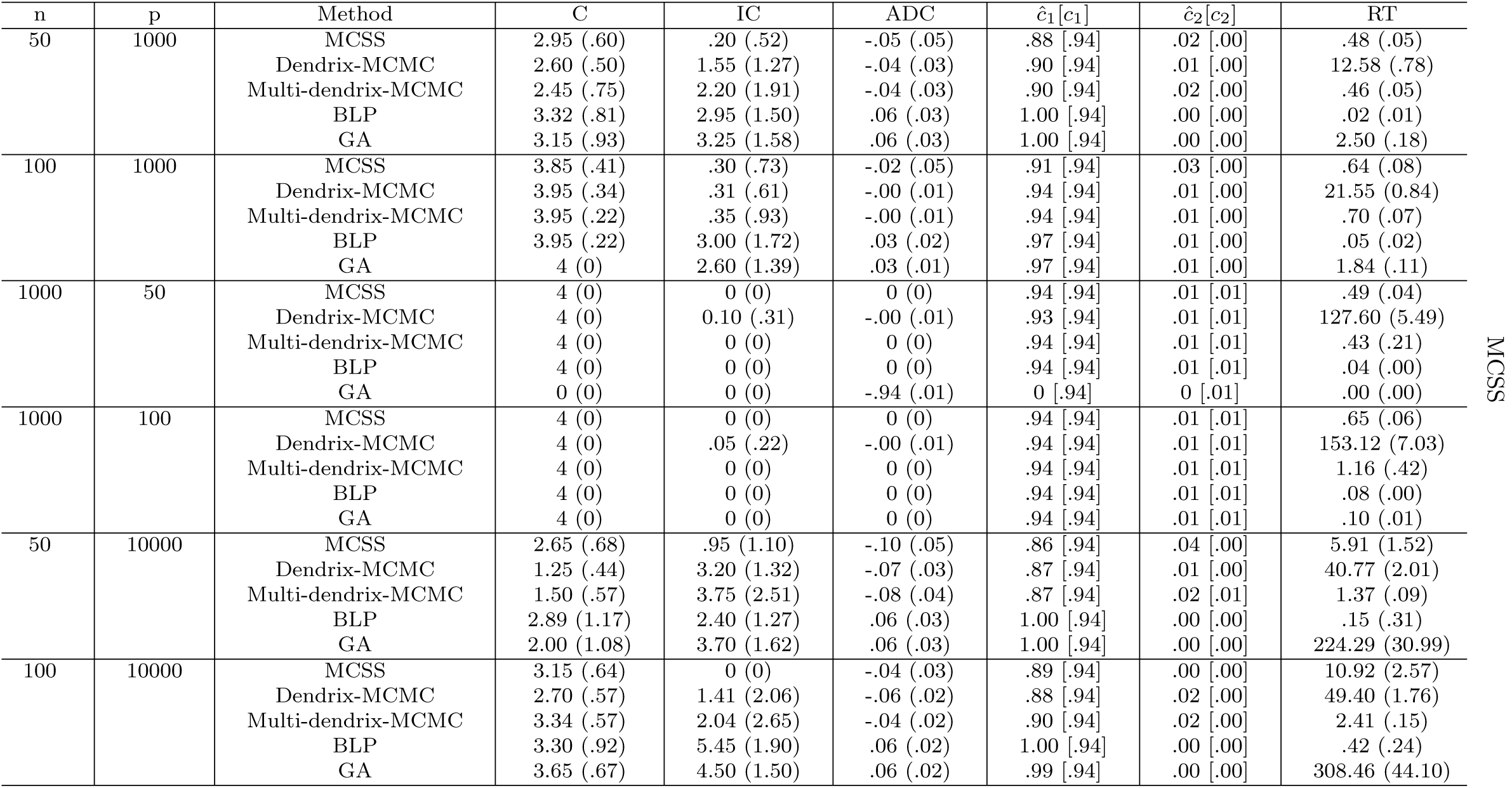
*Results in Simulation I based on 100 simulation replicates with for* (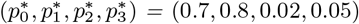).

To evaluate the performance involving cross-validation, consider the first set-up: *n* = 50 and *p* = 1000 with (*p*_1_,*p*_2_,*p*_3_) = (0.95, 0.01, 0.05). The crossvalidation procedure was applied with an enlarged size of Λ, say 100, and Algorithm 1 was applied to *A* for each λ ∈ Λ separately. The results are displayed in Figure 3, demonstrating that the λ’s minimizing the tuning error corresponded to the minimum cost of (2.1) and the true size of *B*_0_, say 4.

**FIG 3.**
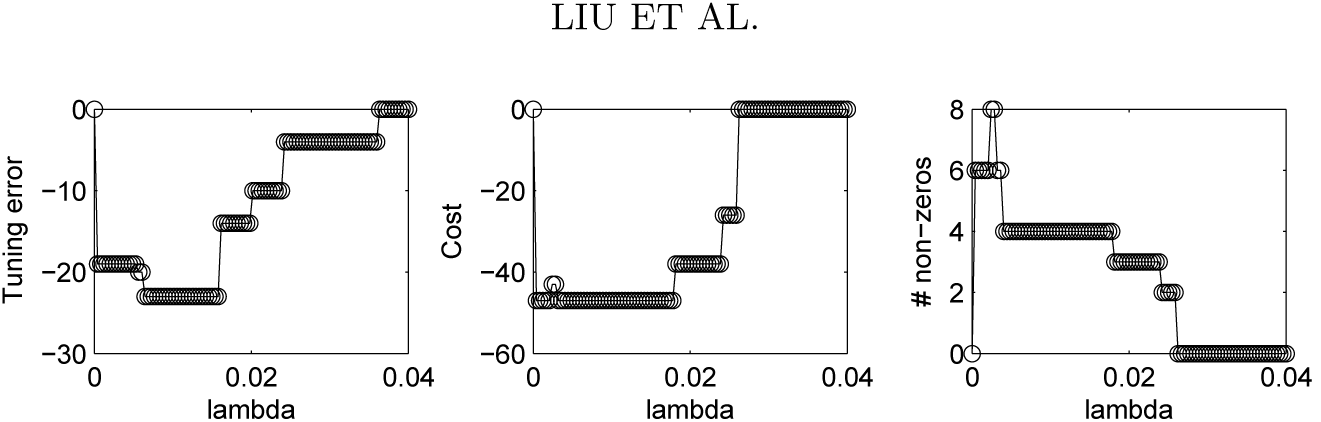
Tuning error, cost and number of non-zero (i.e. true positive) estimates of MCSS versus the tuning parameter value λ ∈ Γ with |Γ|= 100 for the first simulation set-up: *n* = 50, *p* = and (*p*1, *p*2,*p*2) = (0.95, 0.01,0.05).

Moreover, the current tuning error is obtained by applying the crossvalidation procedure for once in consideration of computational efficiency.

For instance, in the first set-up with (*n =* 50, *p* = 1000) and (*p*_1_,*p*_2_,*p*_3_) *=* (0.95,0.01,0.05), as indicated in Figure 4, as the cross-validation fold number increased, the performance of the proposed method measured by C, IC and ADC did not improve, while RT increased linearly.

**FIG 4.**
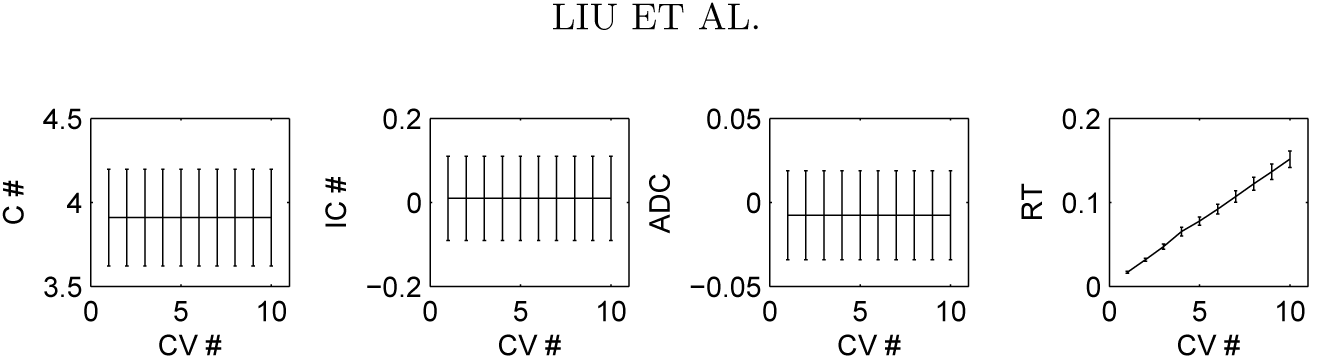
The correct (C#) and incorrect (IC #) numbers of non-zero (i.e. true positive) estimates, the average difference of the costs (ADC) and running time (RT) of MCSS versus the fold number of cross-validation used in Algorithm 2 (CV #) for the first simulation *set-up: n* = 50, *p* ***=*** 1000 *and* (*p*_1_,*p*_2_,*p*_3_) = (0.95, 0.01, 0.05).

#### 3.2.2. Simulation II: multiple driver pathways

We further compared the performance of MCSS against Multi-dendrix in identifying multiple true driver pathways as follows.

The simulation set-up was similar as before except that there were two true driver pathways *B*_1_ and *B*_2_. We used 100 random initial estimates for MCSS. We compared their performance using the top two estimated sets (with the minimum cost function values) by each method for each dataset. As shown in Table 9, MCSS performed much better for the most challenging high-dimensional case with *p* = 10000 and *n* = 50: it correctly identified a much larger number of the genes in the two true driver pathways (i.e. with a larger number of estimated true positives) while yielding fewer false positives. On the other hand, as the sample size *n* increased to 100, the performance of Multi-dendrix caught up.

**TABLE 9.**
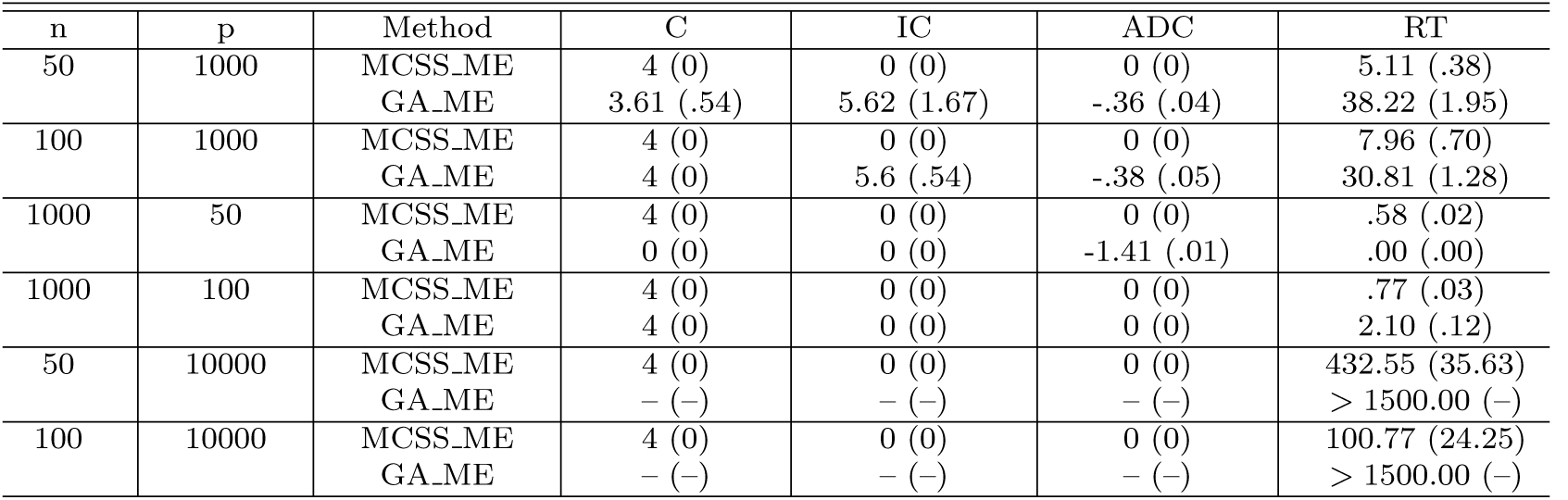
Results in Simulation II based on 100 simulation replications with (*p*_1_,*p*_2_,*p*_3_) = (0.8, 0.02, 0.05).

#### 3.2.3. Simulation III: with both mutation and gene expression data

We generated the mutation data as in Table 7 and the gene expression data from a multivariate normal distribution *N*(0, *V*). Specifically, we divided the genes {1,⋯, *p*} into mutually disjoint subsets *B*_0_ = {1,2,3,4}, *B*_1_, *B*_2_, *…B_K_*, where for each *k* ∈ {1,⋯, *K*}, the gene set size |*B_k_*| was random from {2, ⋯, 20}. V is a correlation matrix with all diagonal elements *Vjj* = 1; for any *j*_1_< *j*_2_ both in the same *B_k_*, 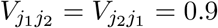; otherwise, 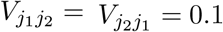. The rationale is that, for the genes in the same set, due to their shared function, their expression levels are also highly correlated. We used our proposed method to select all the tuning parameters, including γ. The simulation results for the integrative analysis of both mutation data and gene expression data are summarized in Table 10, where the new method MCSS_ME is compared with GAME, the integrative version of GA (Zhao *et al.,* 2012). Note that, to our knowledge, the integrative version of BLP in Zhao *et al.* (2012)is not yet publicly available. From Table 10, we see that GA_ME failed in situations with the dimension *p* much smaller than the sample size *n;* in contrast, the new method MCSS_ME performed well. Furthermore, GA_ME was much time-consuming for large *p*.

**TABLE 10.**
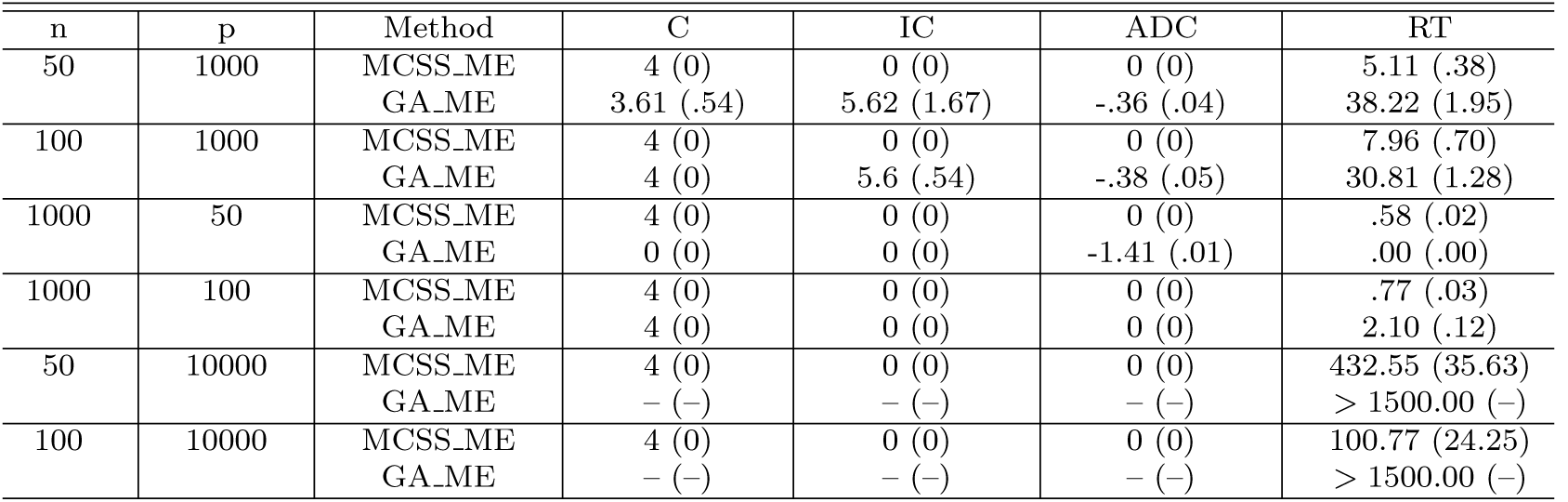
*Results in Simulation III for integrative analysis of mutation data and gene expression data.* (*p*_1_,*p*_2_,*p*_3_) = (0.8,0.02,0.05)

## 4. Conclusions

This paper has introduced a new computational method for a combinatorial optimization problem motivated from cancer genomics. It approximates a combinatorial cost function with a continuous and non-convex relaxation. In particular, the indicator function is approximated by a non-convex truncated *L*_1_-function. The proposed method is computationally more efficient than an existing approach based on stochastic search, and compares favorably over several existing methods in simulations. Through both real data and simulated data analyses, the proposed method was shown to be promising for discovering mutated driver pathways with tumor sequencing data. In light of that Dendrix and other methods have been successfully applied to the TCGA (Kandoth *et al.,* 2013), it would be interesting to apply our proposed method to on-going large cancer genomics projects. Furthermore, the current problem differs from existing pathway analysis of genome-wide association studies (GWAS) (Wang *et al.,* 2007; *Torkamani et al.,* 2007; Schaid *et al.,* 2012) in two aspects: (i) the current problem is more challenging in the sense that no pathway is given a priori; (ii) however, GWAS data is different with genetic variants (or mutations) present for healthy control subjects, and it is also higher-dimensional with a larger number of genetic variants. It would be interesting to see whether the key concept of mutation exclusivity and associated methodology in the current context can be extended and applied to GWAS for de novo pathway or gene subnetwork (Liu *et al.,* 2014) discovery to handle genetic heterogeneity. Finally, the main idea of our algorithm is quite general and may be modified and extended for other challenging combinatorial search problems.

Matlab code implementing the new method and a manual are available at https://github.com/ChongWu-Biostat/MCSS.

## APPENDIX

### Proof of Theorem 1

For convergence of Algorithm 1, by construction, we have, for *m* ∈ ℕ, *S*(*β̂*^(*m*)^) = *S*^(*m+1*)^( *β̂*^(*m*)^) *≤ S*^(*m*)^(*β̂*^(*m*)^) ≤ *S*^(*m*)^(*β̂*^(*m–*1)^) = *S*(*β̂*^(*m–*1)^). Since *S*(*β*) is obviously bounded below, the convergence is proved. Converging finitely follows from the strict decreasing character of *S*^(*m*)^(*β̂*^(*m*)^) in *m*, uniqueness of minimizer of *S*^(*m*)^(*β*) and finite possible values of ∇*S*_2_(*β̂*^(*m–*1)^) in (2.4). After termination occurs at m*, *β̂*^(*m*)^ remains unchanged for *m* ≥ *m*\*, so does the cost function S(*β̂*^(*m*)^) in (2.3) for *m* ≥ *m*\*. By construction of *S*(*β*), we have that *β̃* = *β̂*^(*m*)^ = *β̂*^(*m*–*1)^, for all *m* ≥ *m*\*. *β̃* is uniquely defined, because for each *m* ∈ ℕ, the minimizer *β̂*^(*m*)^ of *S*^(*m*)^ (*β*) is uniquely defined. Since 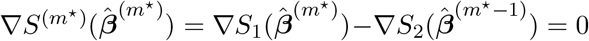, we get that 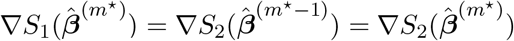. Thus, 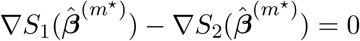, which completes the proof.

### Proof of Lemma 1

We prove by contradiction. By construction of *S*(*β*), we see that 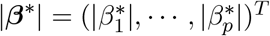 is also a local minimum of *S*(*β*), *β* ∈ *R^p^.* Without loss of generality, we assume that 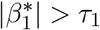. Let

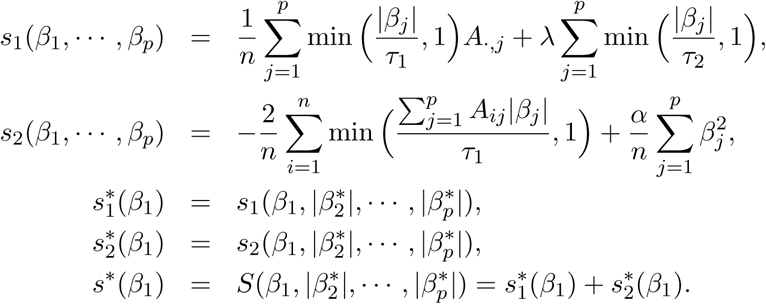

Since (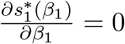 and 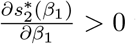 whenever 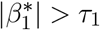 we see that 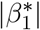 is not a local minimizer of 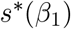, which is contrary to the assumption.

### Proof of Lemma 2

We prove by contradiction. We assume that *β*\* ≠ 0 is a local minimizer of *S*(*β*) in (A.1) on ℝ^*p*^. By construction of *S*(*β*), we see that 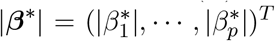
 is also a local minimum of *S*(*β*), *β* ∈ *R^p^*. Without loss of generality, we assume that 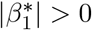. Let

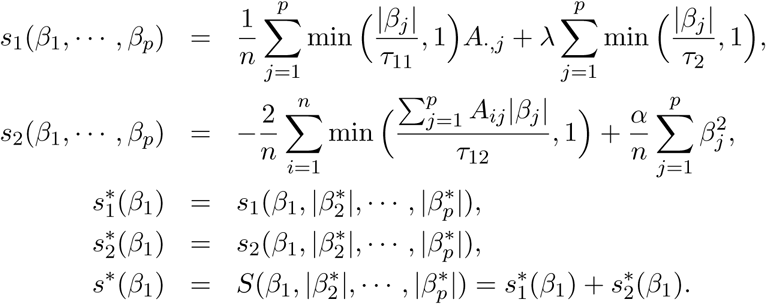

We first consider the situation of 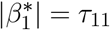. Denote by the right derivative of 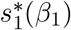 at 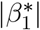 to be b. By construction of 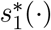, its left derivative at 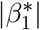 must be 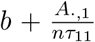. Let *c*_1_ and *c*_2_ denote the left derivative and right derivative of 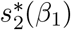 at 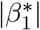 respectively. Since 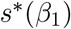 achieves a minimum at 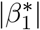, we have that 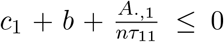 and *c*_2_ + *b* ≥ 0, which implies that 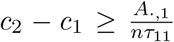. On the other hand, since 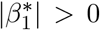, we have that *c*_1_, 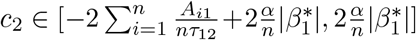, and thus 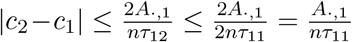 because we have assumed that *τ*_12_ *> 2τ*_11_, which is contrary to the fact that 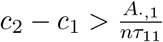.

Second, we consider the situation of 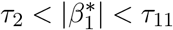. In this situation, the left derivative of 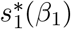 at 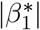, *b*, is 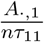, and the left derivative of 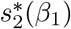 at 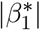, *c*_1_, belongs to 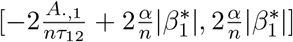, which implies *b* + *c*_1_ > 0 and is contrary to the the assumption of local minimum of 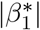.

Third, we consider the situation of 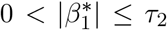. We see that the left derivative of 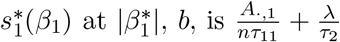, and the left derivative of 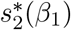 at 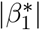, *c*_1_, belongs to 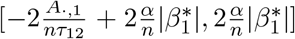, which implies *b* + *c*_1_ > 0 and is contrary to the the assumption of local minimum of 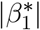.

Finally, we consider the situation of 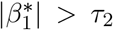. Since 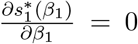 and 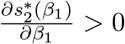 whenever 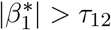, we see that 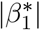 is not a local minimizer of 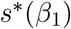, which is contrary to the assumption.

### Other choices of the tuning parameters

This section focuses on situations involving different thresholding parameters for different approximations of indicator functions in (2.3). Consider, for *β* ∈ [0, +∞)^*p*^,

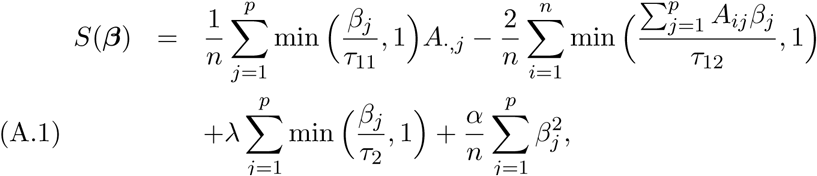
where *τ*_11_ and *τ*_12_ may not be equal.

First, we examine the cases of *τ*_12_ > 2*τ*_11_ (i_2_ < *τ*_11_, *τ*_12_).

#### LEMMA 2.

Let *τ*_12_ *≥* 2*τ*_11_ *and τ*_2_ *< τ*_11_,*τ*_12_*. If there exists a local min-imizer β\** ≠ 0 *of S*(*β*) *in* (A.1), *then 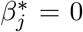 or* 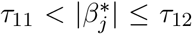 *for each j* ∈ {1, ⋯,*p*}.

Letting *τ*_12_ ≥ 2*τ*_11_, we have that in each iteration of Algorithm 1,

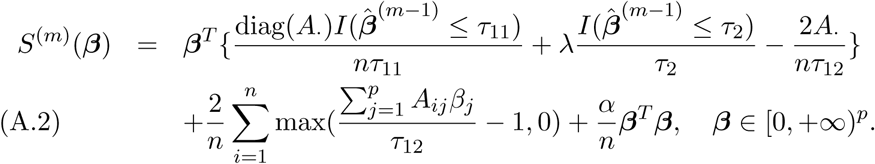

It follows from (A.2) that once we have that *β̂*^(*m–*1)^ = 0 for some *m*, *S*^(*m*)^(*β*) ≥ 0 for all *β* ∈ [0, +∞)^*p*^, which terminates the DC iteration process, because *β̂*^(*m*)^ = *β̂*^(*m–*1)^ = 0. This indicates that if *τ*_12_ ≥ 2*τ*_11_, the DC algorithm becomes sensitive to an initial value *β̂*^(0)^.

Next, we examine the case of 0 < *τ*_12_ < 2*τ*_11_ (*τ*_2_ < *τ*_11_, *τ*_12_), where the DC algorithm is not sensitive as the first one. However, based on the results of a few numerical examples (not shown), we found that in this situation, even using one more parameter, the performance of finding the minimum cost subset did not improve over the proposed method.

Finally, we consider the case of *τ*_1_ = *τ*_11_ = *τ*_12_ and *τ*_2_ ≥ *τ*_1_. In this case, similar to Lemma 1, any local minimizer of *S*(*β*) belongs to [0,*τ*_2_]^*p*^, where the truncated *L*_1_ penalty becomes a *L*_1_ penalty that does not restrict the number of nonzero coordinates of a minimizer as an *L*_0_ penalty does. In particular, in the situation with *τ*_1_ = *τ*_2_, *S*(*β*) becomes a strictly convex function on [0,*τ*_2_]^*p*^, which indicates that for any *β*_1_ and *β*_2_ with 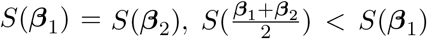. As a result, if there exists two minimum cost subsets *B*_1_ and *B*_2_ in the finite-sample situation, then by using *τ*_1_ = *τ*_2_, the corresponding method is more likely to select *B*_1_ ∪ *B*_2_ as the minimum cost subset.

### The subgradient descent algorithm

For MCSS, we denote 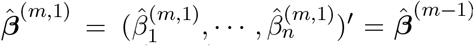, use the following subgradient of *S*^(*m*)^(*β*) at *β̂*^(*m,t–*1)^

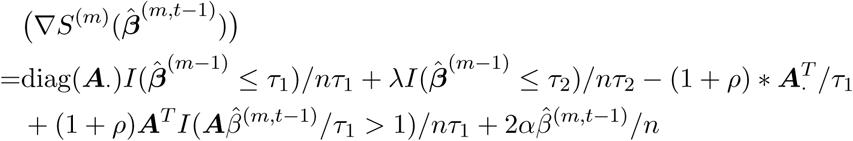

and then update *β̂*^(*m,t*)^ until convergence to obtain *β̂*^(*m*)^:

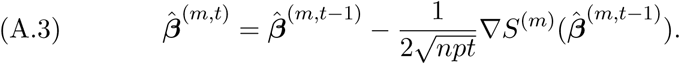

For MCSS_ME, we denote 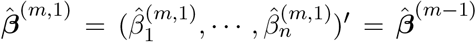, use the following subgradient of *S*^(*m*)^(*β*) at *β̂*^(*m,t–*1)^:

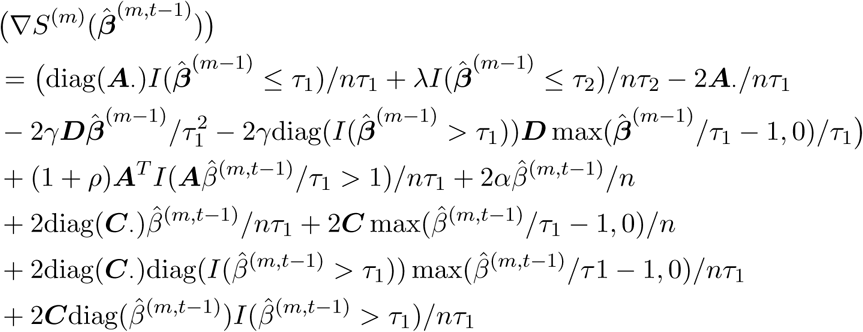

and then update *β̂*^(*m,t*)^ by equation (A.3) until convergence to obtain *β̂*^(*m*)^.

## ACKNOWLEDGMENT

We are grateful to the editors and a reviewer for many constructive and helpful comments. This work was supported by NIH grants R01GM113250, R01HL105397 and R01HL116720, by NSF grants DMS-0906616 and DMS-1207771 and by NSFC grant 11571068. The authors thank Dr. Vandin for sharing the data.

